# Modest static pressure suppresses columnar epithelial cell proliferation in association with cell shape and cytoskeletal modifications

**DOI:** 10.1101/167270

**Authors:** Man Hagiyama, Norikazu Yabuta, Daisuke Okuzaki, Takao Inoue, Yasutoshi Takashima, Ryuichiro Kimura, Aritoshi Ri, Akihiko Ito

## Abstract

Intraluminal pressure elevation can cause degenerative disorders, such as ileus and hydronephrosis, and the threshold is fairly low and constant, 20–30 cm H_2_O. We previously devised a novel two-chamber culture system subjecting cells cultured on a semipermeable membrane to increased culture medium height (water pressure up to 60 cm H_2_O). Here, we cultured several different cell lines using the low static pressure-loadable two-chamber system, and examined cell growth, cell cycle, and cell morphology. Madin–Darby canine kidney (MDCK) columnar epithelial cells were growth-suppressed in a manner dependent on static water pressure ranging from 2–50 cm H_2_O, without cell cycle arrest at any specific phase. Two other types of columnar epithelial cells exhibited similar phenotypes. By contrast, spherical epithelial and mesenchymal cells were not growth-suppressed, even at 50 cm H_2_O. Phalloidin staining revealed that 50 cm H_2_O pressure load vertically flattened and laterally widened columnar epithelial cells and made actin fiber distribution sparse, without affecting total phalloidin intensity per cell. When the mucosal protectant irsogladine maleate (100 nM) was added to 50-cm-high culture medium, MDCK cells were reduced in volume and their doubling time shortened. Cell proliferation and morphology are known to be regulated by the Hippo signaling pathway, but a pressure load of 50 cm H_2_O did not alter the expression levels of Hippo signaling molecules in columnar epithelial cells, suggesting that this pathway was not involved in the pressure-induced phenotypes. RNA sequencing of MDCK cells showed that a 50 cm H_2_O pressure load upregulated *keratin 14*, an intermediate filament, 12-fold. This upregulation was confirmed at the protein level by immunofluorescence, suggesting a role in cytoskeletal reinforcement. These results provide evidence that cell morphology and the cytoskeleton are closely linked to cell growth. Pathological intraluminal pressure elevation may cause mucosal degeneration by acting directly on this linkage.

**Summary:** We provide evidence that columnar epithelial cells are growth-suppressed by pressure loads as low as 30 cm H_2_O, in association with cell-shape flattening and cytoskeletal alterations.

## Introduction

Intraluminal pressure exists at various sites in the body, such as the digestive, biliary, and urinary tracts, and intraocular, intracranial, and intraventricular spaces. The normal mean values range approximately from 5 to 20 cm H_2_O (Bratt and Nilsson, 1987; Coelho, et al., 1985; Enevoldsen et al., 1976; Kawoos et al., 2015; Liu et al., 2003; Schmidt et al., 1978). Increases in intraluminal pressure are thought to cause several diseases and disorders, such as ileus, jaundice, hydronephrosis, hydrocephalus, and glaucoma (Kumar, et al., 2010; Lindahl, et al., 1995). These pathological conditions involve degenerative lesions of the mucosa and nervous system, and are known to emerge when intraluminal pressure is persistently elevated over 20–30 cm H_2_O (Bratt and Nilsson, 1987; Kawoos et al., 2015; Leske et al., 2003). Thus, the threshold for pressure-induced degeneration is fairly low and constant, regardless of the tissues or organs where the pressure is loaded. This suggests that there is a common mechanism by which such low pressure causes cell and/or tissue degeneration, but the precise molecular basis has not yet been intensively examined.

Recently, we devised a new two-chamber culture system, in which small amounts of water pressure can be loaded on cells without disturbing the standard culture conditions, such as normal pH and partial pressures of oxygen and carbon dioxide (Yoneshige et al., in press). Using this system, we demonstrated that mouse superior cervical ganglion neurons degenerated when water pressure was over 30 cm H_2_O (Yoneshige et al., in press). Cell adhesion molecule 1, an immunoglobulin superfamily member, appeared to be involved in this neuronal degeneration through increased ectodomain shedding to produce a C-terminal fragment that accumulates in neurites (Yoneshige et al., in press).

On the other hand, mucosal degeneration associated with intraluminal pressure elevation is thought to involve the retardation of mucosal epithelial renewal, suggesting that pressure directly suppresses epithelial cell proliferation. This speculation is reasonable but its mechanisms have not been carefully examined. The Hippo signaling pathway may be involved because it is crucial for controlling epithelial cell growth by regulating contact inhibition (Gumbiner and Kim, 2014; Yu et al., 2015). This pathway consists of a molecular cascade involving MST2, LATS1, LATS2, YAP, and TAZ (Yu et al., 2015). When the pathway is inactive, YAP is present in the nucleus and transactivates target genes, promoting cell proliferation (Yu et al., 2015). When the pathway is activated, YAP is phosphorylated and retained in the cytoplasm, resulting in suppression of target gene transactivation and cell proliferation (Yu et al., 2015).

Irsogladine maleate (IM) has long been widely used as a mucosal protectant that promotes the healing of gastric ulcers, based on its ability to reinforce epithelial cell adhesion by increasing adhesion-related molecule expression, gap junctional cell–cell communication, and cAMP production (Akagi et al., 2013). Thus, IM may counteract the action of water pressure on epithelial cells.

In the present study, we cultured various types of epithelial and mesenchymal cells using a water pressure-loadable two-chamber system, and examined changes in cell growth profiles and cell morphology. Next, we analyzed protein expression of the Hippo pathway molecules and comprehensively compared gene expression between pressure-loaded and non-loaded epithelial cells by RNA sequencing. In addition, we examined whether IM rescued the pressure-induced phenomena of epithelial cells. Pressure-induced phenotypes revealed a close link among morphology, cytoskeleton, and proliferation in columnar epithelial cells.

## Results

### Low static pressure flattens columnar epithelial cells and suppresses their proliferation

Madin–Darby canine kidney (MDCK) cells, which are of epithelial origin with a columnar morphology, were cultured on a semipermeable membrane below culture medium that was 2, 15, 30, or 50 cm in height, and the cell numbers were counted after 1, 2, and 3 days. The cell growth curve showed a slow ascent and the doubling time lengthened in a culture medium height-dependent manner (Fig. 1A). Two other columnar epithelial cell lines, NCI-H441 and Caco-2, and two mesenchymal cell lines, TIG-1 and NIH3T3, were cultured in 2- or 50-cm-high culture medium. NCI-H441 and Caco-2 cells produced results similar to those of MDCK cells (Fig. S1A, D), whereas TIG1 and NIH3T3 cells did not exhibit significant differences between the two culture conditions (Fig. S2A, C). KATO-III and NUGC-4 cells, which are of epithelial origin but have a spherical morphology, were not growth-suppressed by a static pressure of 50 cm H_2_O (Fig. S3A, C). Cell cycle profiles of MDCK, NCI-H441 and Caco-2 cells were analyzed by flow cytometry. There were no apparent growth-arrested phases detectable in cells in 50-cm-high-medium cultures (Fig. 1B; Fig. S1B, E).

**Figure 1.**
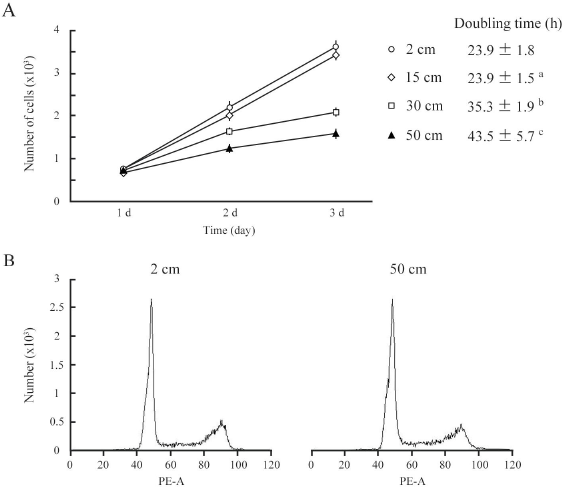
Low static pressure suppresses MDCK cell proliferation. A. MDCK cells were cultured on a semipermeable membrane in 2-, 15-, 30-, or 50-cm-high culture medium for 3 days. Cell numbers were counted every day, and were plotted with bars indicating standard deviations. Cell doubling time was calculated. ^a^, ^b^, and ^c^, *P* = 0.985, < 0.001, and < 0.001, respectively, by Student’s *t*-test when compared with 2-cm-high-medium cultures. B. After 3 days of culture in 2- or 50-cm-high medium, MDCK cells were labeled with propidium iodide and analyzed by flow cytometry. Representative results are shown.

Cultured cells were stained with phalloidin. MDCK cells were found to change drastically in shape with the application of a static pressure of 50 cm H_2_O. The cell area tripled in the XY plane and the cell height roughly halved in the Z-axis, resulting in 1.5-fold enlargement of the cell volume (cell area × cell height) (Fig. 2). Phalloidin staining was markedly weakened in the cytoplasm and on the cell membrane when XY planes around the middle of the Z-axis were observed, but the total intensity per cell was unchanged when the cell volume increase was taken into consideration (Fig. 2), suggesting that actin fibers became sparsely distributed as the cell volume increased. Similar results were obtained with NCI-H441 and Caco-2 cells (Fig. S1C, F). By contrast, in 50-cm-high-medium cultures of KATO-III and NUGC-4 cells, some cells showed a little flattened morphology, but there was no apparent decrease in phalloidin staining intensities (Fig. S3B, D). Neither TIG-1 nor NIH3T3 cells exhibited substantial differences in cell shape or phalloidin staining intensity between 2- and 50-cm-high-medium cultures (Fig. S2B, D).

**Figure 2.**
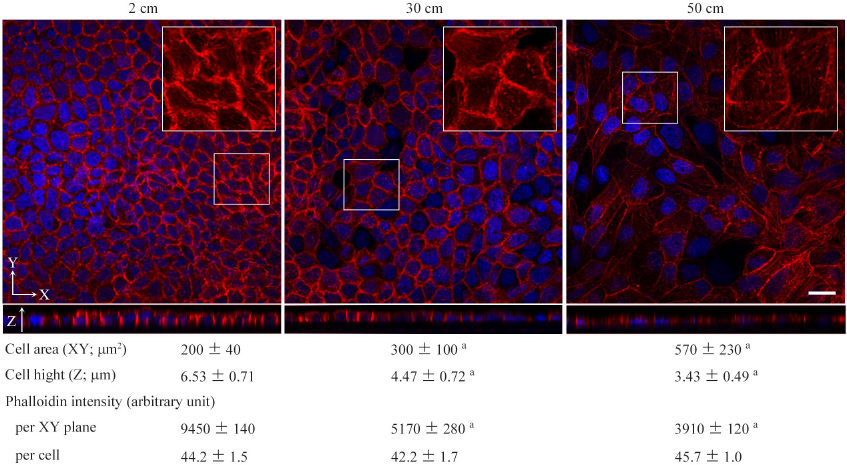
Low static pressure flattens epithelial cells in association with diffusion of phalloidin staining. MDCK cells were cultured on a semipermeable membrane in 2-, 30-, or 50-cm-high culture medium for 3 days, and stained with a mixture of phalloidin (red) and DAPI (blue). Cell area and height and phalloidin intensity were measured by laser microscopic examinations. The total intensities of phalloidin per XY plane and per cell were calculated. The former is the mean intensity of three XY planes around the middle of the Z-axis. The latter is calculated as follows: the mean phalloidin intensity of ten randomly selected ZX planes was multiplied by the area of the XY plane, then divided by the cell number. Boxed areas are enlarged to depict intracellular phalloidin staining, where DAPI fluorescence is not merged. ^a^ *P* ≤ 0.001 by Student’s *t*-test when compared with 2-cm-high-medium cultures. Scale bar = 20 μm.

### Low static pressure does not alter protein expression of Hippo pathway molecules

MDCK cells were cultured in 2- or 50-cm-high culture medium for 3 days, and their protein extracts were fractionated into cytosolic and nuclear fractions. Successful fractionation was confirmed by western blot analyses with antibodies against glyceraldehyde 3-phosphate dehydrogenase (GAPDH) and lamin B (Fig. 3). The duplicate blots were examined for expression of the Hippo pathway molecules, MST2, LATS1, LATS2, YAP, and TAZ. No appreciable differences were detected in the expression levels of the five molecules between the two culture conditions in either fraction (Fig. 3). Similar results were obtained with NCI-H441 and Caco-2 cells (Fig. 3)

**Figure 3.**
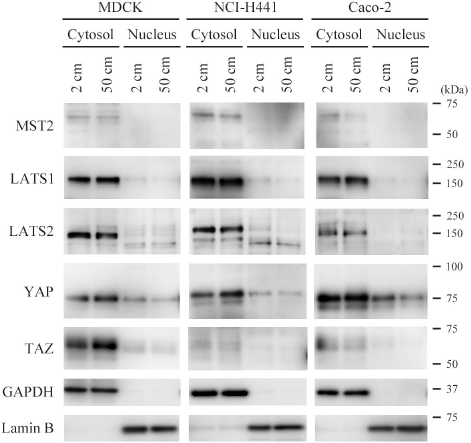
Expression analyses of Hippo pathway molecules. MDCK, NCI-H441, and Caco-2 cells were cultured on a semipermeable membrane in 2- or 50-cm-high medium for 3 days. Cytosolic and nuclear proteins were separately extracted from the cells, and were blotted with the antibodies indicated. Glyceraldehyde 3-phosphate dehydrogenase (GAPDH) and lamin B were used as cytoplasmic and nuclear markers, respectively.

### Upregulation of keratin 14 in MDCK cells by 50 cm H_2_O pressure

RNAs were extracted from MDCK cells cultured in 2- and 50-cm-high culture medium, and gene expression differences were compared by RNA sequencing. The raw data have been deposited in the NCBI Gene Expression Omnibus database (GSE100794). The gene most upregulated in 50-cm-high-medium cultures was *keratin 14*, encoding a type I intermediate filament, with an 11.869-fold change (Fig. 4A). This upregulation was confirmed at the protein level by western blot analyses (Fig. 4B). MDCK cells were double-stained with phalloidin and an anti-keratin 14 antibody. Keratin 14 expression was at trace levels in 2-cm-high-medium cultures, whereas it was clearly detected in the cytoplasm of MDCK cells cultured in 50-cm-high medium, in contrast to a marked decrease in phalloidin staining (Fig. 4C).

**Figure 4.**
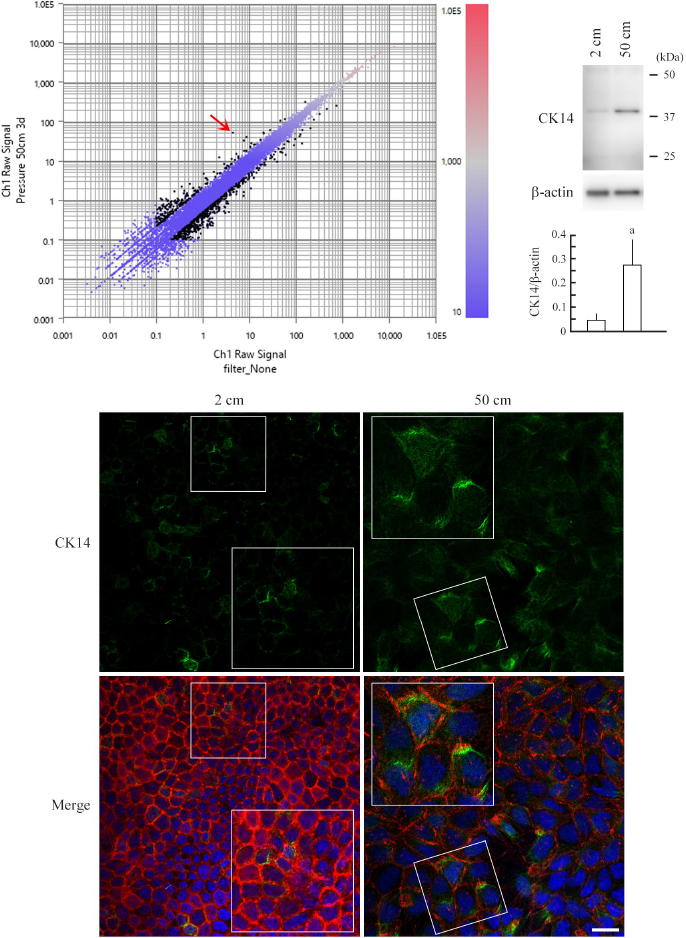
RNA sequencing of MDCK cells and upregulation of keratin 14 in pressure-loaded MDCK cells. A. Total RNA was extracted from MDCK cells that were cultured in 2- or 50-cm-high medium, and was subjected to RNA sequencing-based gene expression comparison analysis. The data are expressed as a scatter plot comparing the log_10_ fragments per kilobase of exon per million mapped fragments (FPKM) values for 2-cm-high-medium cultures (X-axis) versus 50-cm-high-medium cultures (Y-axis). The FPKM values are displayed in a red-to-blue gradation, with blue and red indicating the lowest and highest values, respectively. Black dots indicate genes with > 2-fold change. *Keratin 14*, the gene most upregulated in 50-cm-high-medium cultures, with an 11.869-fold change, is indicated with a red arrow. B. MDCK cells were cultured in 2- or 50-cm-high medium, and were subjected to western blot analysis using an anti-keratin 14 antibody (upper panel). The blot was reprobed with an anti-β-actin antibody to indicate the amount of protein loading per lane. Intensities of the immunoreactive bands for keratin 14 and β-actin were measured densitometrically. The mean ratios of keratin 14 to β-actin and standard deviations were calculated from the data obtained in three independent experiments, and were statistically analyzed by Student’s *t*-test. ^a^ *P* = 0.019 when compared with 2-cm-high-medium cultures. C. MDCK cells were cultured on a semipermeable membrane in 2- or 50-cm-high medium for 3 days, then were triple-stained with keratin 14 immunofluorescence (green; upper), phalloidin labeling (red), and DAPI nuclear staining (blue). The three images are merged (lower). Boxed areas are enlarged to depict keratin 14 upregulation in the cytoplasm of pressure-loaded cells. Scale bar = 20 μm.

### IM significantly rescues pressure-induced phenomena

MDCK cells were cultured on a semipermeable membrane for 3 days in 2- or 50-cm-high culture medium containing either IM or vehicle (dimethyl sulfoxide, DMSO) alone (final concentrations of IM and DMSO, 100 nM and 10,000× dilution, respectively). Cell doubling times in 2-cm-high-medium cultures were comparable in the IM and vehicle groups, whereas in 50-cm-high-medium cultures, the IM group had a significantly shorter doubling time than the vehicle group (Fig. 5A). Cell cycle analyses did not indicate any apparent differences between the two groups (Fig 5B). MDCK cells were stained with phalloidin. Cell morphology and phalloidin staining intensity were comparable between the IM and vehicle groups in 2-cm-high-medium cultures (Fig. 6). When cultured in 50-cm-high medium, the IM group had a smaller XY plane cell area, a larger cell height, and a smaller cell volume than the vehicle group (Fig. 6). IM-treated cells stained more strongly with phalloidin than the vehicle-treated cells, when observed in XY planes around the middle of the Z-axis. The total intensity per cell, however, was comparable in the IM and vehicle groups (Fig. 6), suggesting that IM did not affect the kinetics of actin polymerization/depolymerization.

**Figure 5.**
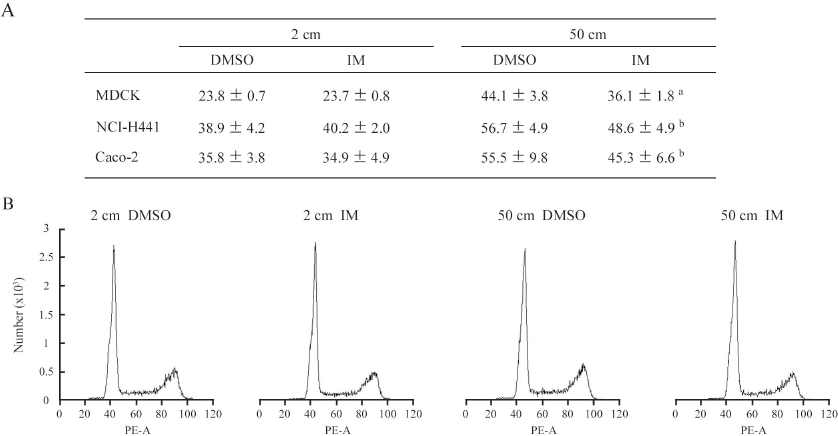
Cell growth and cell cycle analyses in the presence of irsogladine maleate. A. MDCK, NCI-H441, and Caco-2 cells were cultured in 2- or 50-cm-high medium containing either irsogladine maleate (IM; 100 nM) or DMSO (10,000× dilution), and their doubling times were calculated. The mean and standard deviation (hour) are shown. a and b, *P* ≤ 0.01 and 0.05, respectively, by Student’s *t*-test when compared with the 50 cm DMSO group. B. After 3 days of culture in 2- or 50-cm-high medium, MDCK cells were labeled with propidium iodide and analyzed by flow cytometry. Representative results are shown.

**Figure 6.**
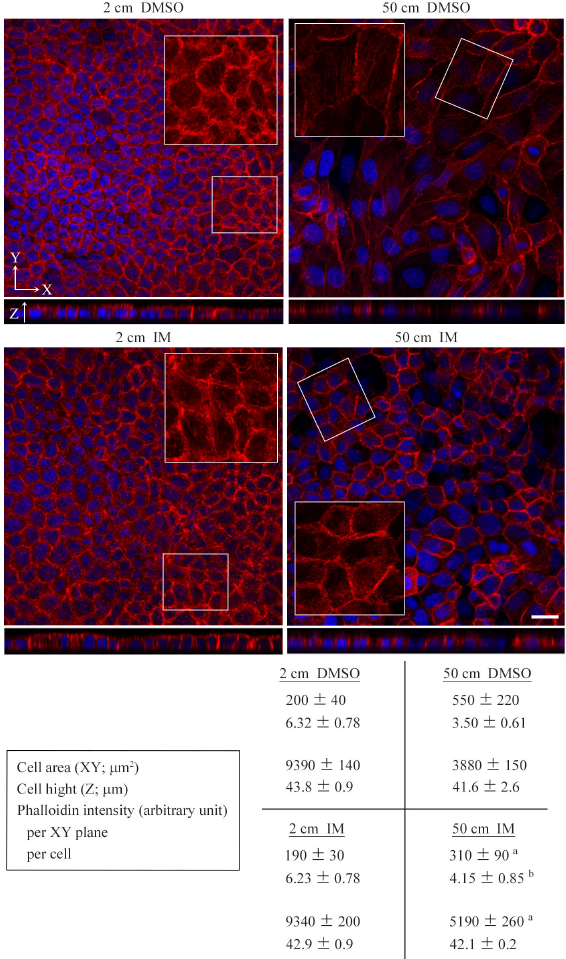
Effects of irsogladine maleate on cell morphology and phalloidin staining. MDCK cells were cultured on a semipermeable membrane for 3 days in 2- or 50-cm-high medium containing either irsogladine maleate (IM; 100 nM) or DMSO (10,000× dilution), and stained with a mixture of phalloidin (red) and DAPI (blue). Cell area and height and phalloidin intensity were measured by laser microscopic examinations. The total intensities of phalloidin per XY plane and cell were calculated. The former is the mean intensity of three XY planes around the middle of the Z-axis. The latter is caluculated as follows: the mean phalloidin intensity of 10 randomly selected ZX planes was multiplied by the area of the XY plane, then divided by the cell number. Boxed areas are enlarged to depict intracellular phalloidin staining, where DAPI fluorescence is not merged. ^a^ and b, *P* ≤ 0.01 and 0.05, respectively, by Student’s *t*-test when compared with the 50 cm DMSO group. Scale bar = 20 μm.

### Discussion

In the present study, we found that relatively small increases in water pressure significantly suppressed the growth of columnar epithelial cells in association with cell height shortening and lateral widening. This cell shape change was associated with decreased intensity of phalloidin staining in the XY plane. By contrast, spherical epithelial-origin cells and mesenchymal cells were not growth-suppressed by the pressure load of 50 cm H_2_O, and did not exhibit noticeable changes in cell morphology or phalloidin staining. These results suggest that cell morphological changes and cytoskeletal remodeling are involved in growth suppression induced by modest static pressure, and are consistent with the notion that actin fibers act as a mechano-sensing apparatus (Hayakawa et al., 2011). A pressure of 50 cm H_2_O is nearly equal to 0.05 atm = 5 kPa = 5 fN nm^−2^. Nanometer-sized molecules in live cells fluctuate with forces on the piconewton order because the thermal energy of each molecule at room temperature is on the order of multiplying nanometers by piconewtons (*k*_B_*T* = 4.1×10^−21^ J = 4.1 pN·nm) (Merkel et al., 1999). This means that under a pressure of 5 fN nm^−2^, the force loaded on each molecule is fN order, and is too small to perturb the molecular fluctuation. How can cells convert this weak force into biological signals? According to our present study, the primary event caused by the pressure appeared to be cell-shape change, a kind of physical response. This event may trigger particular biological responses, including growth suppression and gene transactivation.

A pressure of 50 cm H_2_O made columnar epithelial cells 1.5-fold larger in cell volume, with lateral widening and cell height shortening. This cell volume increase is likely associated with an increase in cell membrane tension. Actin fibers appeared to become sparsely distributed as a result of cell volume increase because their total amount per cell was unchanged. At this time, individual actin fibers may be stretched out, as they have an extensibility as high as 200% (Labouesse et al., 2016). These events may be involved in pressure-induced cell growth retardation. Increased tension of the cell membrane may interfere with the progression of the cell cycle into the S phase because in cells proceeding to cell division, cell volume gradually increases but cell membrane tension remains low (Chang et al., 2014). Sparse distribution and/or stretching of actin fibers may do the same because preexisting actin filaments are used preferentially for the formation of the contractile ring in dividing cells (Cao and Wang, 1990). These notions are supported by the results obtained from adding IM to cultures: when 50 cm H_2_O pressure-loaded MDCK cells were treated with IM, their doubling time and cell volume decreased significantly, and phalloidin staining strengthened in the XY planes. Although these speculations require further evaluation, the present study provides evidence that cell shape and cytoskeleton play key roles in cell cycle progression.

Notably, pressure-induced cell growth retardation was not associated with cell cycle arrest at particular phases, indicating that low pressure does not activate cell cycle checkpoint systems. Additionally, the Hippo pathway did not appear to be involved in cell growth retardation. This seems reasonable, since the Hippo pathway is known to activate by sensing the failure of actin polymerization (Reddy et al., 2013), but the amounts of actin fibers were comparable between pressure-loaded and non-loaded cells, as discussed above; that is, pressure load appeared to have little influence on the kinetics of actin polymerization/depolymerization. Cell growth retardation may be simply due to a general suppression of metabolism and decline in molecular functions within the pressure-loaded cell. These conditions appear similar to those under starvation. Future studies should clarify the differences between low pressure load and starvation, which induces cell cycle arrest at the G1 phase (Cooper, 2003).

RNA sequencing indicated that *keratin 14* was upregulated by modest static pressure in MDCK cells. This upregulation is reasonable because intermediate filaments including keratin 14 act to reinforce cell stiffness and to resist physical strain imposed on the cell (Ma et al., 1999; Nolting et al., 2014). Expression levels of keratin 14 are shown to change dynamically among cell cycle phases, and this change is apparently required for the cell division process to progress (Confalonieri et al., 2017). Constant upregulation of this keratin in MDCK cells can be regarded as one of the causes of cell growth suppression induced by pressure loading.

Intraluminal pressure elevation often causes the formation of tissue degenerative lesions. Hydronephrosis/obstructive nephropathy leads to renal tubular injury through urinary intraluminal pressure increase (Bratt and Nilsson, 1987; Kipari et al., 2006). Obstruction of pancreatic and biliary ducts results in ductal dilatation through intraductal pressure elevation (Turowski et al., 2011). Mechanical ventilation often leads to alveolar damage through excess airway pressure (Cabrera-Benitez et al., 2012). Delayed tissue regeneration and re-epithelialization are assumed to underlie the development of these lesions (Cabrera-Benitez et al., 2012; Kipari et al., 2006; Turowski et al., 2011). Intraluminal pressure elevation can be a direct cause of delayed tissue regeneration/re-epithelialization through the suppression of epithelial cell proliferation. The present study identified IM as a potent drug that can protect against degenerative lesion formation.

In conclusion, the growth of columnar epithelial cells appeared to be suppressed by a static pressure burden as low as a few tens of cm H_2_O. The precise mechanisms by which cells can sense such modest pressure remain unknown; however, this study highlights a causative role for the cytoskeleton and cell morphology in the proliferative responses of epithelial cells to modest pressure. This linkage may underlie a general mechanism of the development and progression of columnar epithelial degeneration under pathological conditions of intraluminal pressure elevation.

## Materials and Methods

### Cells, antibodies, and reagents

MDCK, NIH3T3, and TIG-1 cells were purchased and cultured as described in our previous reports. NCI-H441 cells (lot no. 58294188) were purchased from the American Type Culture Collection (Manassas, VA, USA) and grown as previously described. Caco-2, KATO-III, and NUGC-4 cells were purchased from the Riken BioResource Center, Tsukuba, Japan. All experiments using these cells were performed within four months after resuscitation.

Primary antibodies used in this study targeted MST2 (#3952; Cell Signaling, Beverly, MA, USA), LATS1 (C66B5; Cell Signaling), LATS2 (#A300-479A, Bethyl Laboratories, Montgomery, TX, USA), YAP (#4912; Cell Signaling), TAZ (#HPA007415; Sigma-Aldrich, St. Louis, MO, USA), keratin 14 (LL002; Dako, Glostrup, Denmark), lamin B (M-20; Santa Cruz, Dallas, TX, USA), β-actin (Medical & Biological Laboratories, Nagoya, Japan), and GAPDH (Medical & Biological Laboratories). Peroxidase-conjugated secondary antibodies used for western blot analysis were purchased from Amersham (Buckinghamshire, England). Phalloidin (rhodamine conjugated) and DAPI were purchased from Molecular Probes (Carlsbad, CA, USA) and Dojindo (Kumamoto, Japan), respectively. IM was kindly provided by Nippon Shinyaku Co., Ltd. (Kyoto, Japan), and was dissolved in DMSO at a concentration of 1 mM (stock solution).

### Two-chamber culture system for water pressure loading

The water pressure-loadable two-chamber culture device was previously described in detail (Yoneshige et al., in press). Briefly, the upper chamber composite consisted of a long plastic cylinder with a water-tight connection with a culture insert lined with a semipermeable membrane, and the unit was placed vertically in a 10-cm dish lower chamber. Between the two chambers, a porous (150 μm, 200 cm^2^) silicon sheet was inserted to support the semipermeable membrane against the medium (water pressure) applied to the upper chamber cylinder. Using this device, cells were subjected to water pressure levels (cm H_2_O) dictated by the height of the medium from the surface to the semipermeable membrane. Partial pressures of oxygen and carbon dioxide and pH were confirmed to be comparable in the upper and lower chambers (Yoneshige et al., in press).

4 × 10^4^ MECK or Caco-2 cells, 8 × 10^4^ NCI-H441 cells, and 1 × 10^5^ NIH3T3, TIG-1, KATO-III, or NUGC-4 cells were seeded onto the bottom of a culture insert lined with a semipermeable membrane (transparent PET membrane, pore size 1.0 μm; Corning Life Sciences, Durham, NC, USA) placed in a 6-well plate, and cells were grown in 2-cm-high Eagle’s minimum essential medium (EMEM, for MDCK, Caco-2, and TIG-1), RPMI1640 (for NCI-H441, KATO-III, and NUGC-4), or Dulbecco’s modified Eagle’s medium (DMEM, for NIH3T3) (Wako Pure Chemical Industries, Osaka, Japan) containing 10% fetal bovine serum (FBS). The following day, when the cells grew to approximately 40–50% confluency, the insert was set in the pressure-loadable two-chamber system, and the upper and lower chambers were filled with the appropriate medium containing 10% FBS. In some experiments, IM stock solution or DMSO was added to the upper chamber at 10,000× dilution.

Cell doubling time was calculated as described previously (Nakamoto et al., 2001). Cell cycle analysis was performed using propidium iodide and flow cytometry, as described previously (Ito et al., 2000). Experiments were done in triplicate, with essentially similar results.

### RNA sequencing

Total RNA was extracted from MDCK cells cultured in 2- and 50-cm-high media using a miRNeasy Mini Kit (Qiagen, Valencia, CA, USA) according to the manufacturer’s protocol. Library preparation was performed using a TruSeq stranded mRNA sample prep kit (Illumina, San Diego, CA, USA) according to the manufacturer’s instructions. Sequencing was performed on an Illumina HiSeq 2500 platform in 75-base single-end mode. Illumina Casava 1.8.2 software was used for base-calling. Sequenced reads were mapped to the dog reference genome (CanFam3.1) using TopHat v. 2.0.13 in combination with Bowtie 2 v. 2.2.3 and SAMtools v. 0.1.19. The fragments per kilobase of exon per million mapped fragments (FPKMs) was calculated using Cuffnorm v. 2.2.1. Among the genes calculated with a normalized FPKM value greater than 1.0 in both MDCK cell cultures, 730 genes were more than two-fold up- or downregulated from the 2-cm H_2_O to 50-cm H_2_O pressure load.

### Protein extraction and western blot analysis

Protein extraction from cultured cells and western blot analysis were performed as described previously (Koma et al., 2008). Immunoreactive band intensities were quantified using ImageJ software (National Institutes of Health, Bethesda, MD, USA), as described previously (Mimae et al., 2011).

### Fluorescent staining and confocal microscopy

Phalloidin staining and nuclear labeling with DAPI were performed as previously described (Ito et al., 2012; Mimae et al., 2011). Immunofluorescence coupled with phalloidin labeling was performed as previously described (Ito et al., 2012). Briefly, cells were fixed in paraformaldehyde, blocked with 2% bovine serum albumin, and incubated with an antibody against keratin 14, then visualized with Alexa Flour 488-conjugated secondary antibody. After washing three times with phosphate buffered saline, phalloidin staining was performed for 10 h at 4°C. Nuclei were labeled with DAPI. Fluorescence images were captured using a C2+ confocal scanning system equipped with 488-nm argon and 543-nm helium–neon lasers (Nikon, Tokyo, Japan). The vertical sectional (Z-plane) images were generated by Z-stack confocal microscopy using a 0.4-μm motor step. Captured images were analyzed on the Nikon C2+ computer system using Analysis Controls tools. Length and area were measured with Annotations and Measurements, cell numbers were counted as the number of DAPI-stained nuclei identified using Object Count, and phalloidin intensity (arbitrary unit) was calculated using ROI Statistics. The mean total phalloidin intensity per cell was calculated as follows: in a high-power field view using a 60× objective lens, the total intensities were measured in three XY planes that were in the middle of the Z-axis, and 0.8 μm up and 0.8 μm down, and the mean value was calculated and expressed as phalloidin intensity per XY plane. The total intensity of phalloidin per cell was calculated as follows: the mean phalloidin intensity of 10 randomly selected ZX planes was multiplied by the area of the XY plane, then divided by the cell number. Ten randomly selected high-power field views were examined per experimental group, and the mean and standard deviation were calculated.

## Statistical analysis

Differences among experimental groups were analyzed using Student’s *t*-test for quantification. A *P*-value ≤ 0.05 was considered to indicate statistical significance.

## Author contributions

MH carried out the cell culture, cell staining, western blotting, and laser microscopic examinations, and performed the statistical analyses. TI, YT, RK, and AR participated in the western blot analysis and microscopic examinations. NY supervised the western blot analysis and data interpretation. DO carried out RNA sequencing and deposited the data into the public database. AI conceived and designed the study, and drafted the manuscript. All authors read and approved the final manuscript.

## Data availability

The raw data of RNA sequencing are deposited in the NCBI Gene Expression Omnibus database (accession number, GSE100794). The URL is as follows. https://www.ncbi.nlm.nih.gov/geo/query/acc.cgi?acc=GSE100794

